# Neuromuscular and biomechanical functions subserving finger dexterity in musicians

**DOI:** 10.1101/625558

**Authors:** Yudai Kimoto, Takanori Oku, Shinichi Furuya

## Abstract

Exceptional finger dexterity enables skillful motor actions such as those required for musical performance. However, it has been not known whether specialized neuromuscular or biomechanical features subserve the dexterity. We aimed to identify the features firstly differentiating the finger dexterity between trained and untrained individuals and secondly accounting for the individual differences in the dexterity across trained individuals. To this aim, two studies were conducted. The first study (Study 1) compared the finger motor dexterity and several neuromuscular and biomechanical characteristics of the fingers between pianists and musically untrained individuals (non-musicians). The results showed no differences in any biomechanical features between the groups with different musical expertise. However, the pianists exhibited faster individual finger movements and more independent control of movements between the fingers. These observations indicate expertise-dependence of the finger motor skills and neuromuscular control of the fingers. The second study (Study 2) assessed individual differences in the finger dexterity between trained pianists. A penalized regression determined an association of the maximum movement speed of the pianists’ individual fingers with both finger muscular strength and biomechanical characteristics of the hand, but not with independent movement control between the fingers. In addition, none of these features covaried with measures of early and deliberate piano practice. Taken together, these findings indicate that distinct biological factors of finger motor dexterity differentiate between the effects of piano training and individual differences across skilled pianists.

## INTRODUCTION

Ubiquitous features of the hand provide dexterity that enables skillful manipulation of various tools, an ability that is necessary for medical procedures, manufacturing, and performing arts such as musical performance. A number of studies have attempted to identify neural and biomechanical attributes that play roles in producing dexterous finger movements ^1^, which includes neuromuscular functions ^2^, inhibitory and excitatory functions of motor cortical neurons ^3–5^, and the anatomical architecture of the hand ^6–8^. Most of these biological features can be adapted through training ^9^. However, this sometimes results in maladaptive changes in the sensorimotor system and causes movement disorders such as focal dystonia ^10,11^. In addition, many movement disorders, such as stroke, cerebellar dysfunctions, and carpal tunnel syndrome, degrades manual dexterity, which severely impairs the quality of life ^12,13^. To shed lights on neural and biomechanical mechanisms subserving the motor dexterity of the fingers is inevitable for optimizing motor training and rehabilitation.

Independent motor control between the fingers plays a key role in dexterity of the hand ^14–17^. The inter-finger independence allows for the simultaneous production of multiple motor actions by the individual fingers, such as moving one finger while doing the preparatory motion of another finger during a sequential finger movement. This motor skill thus provides a basis of variation of movement repertories ^18^. The independent finger motor control is determined by neuromuscular and biomechanical constraints on the fingers ^19^. The neuromuscular constraints can be the synchronous firing of motor neurons innervating into multiple compartments of the same finger muscle ^20–22^ and the shared representation of individual fingers in the motor cortex ^23,24^. Surround inhibition is a mechanism that inhibits the excitability of motor cortical neurons innervating into a finger adjacent to a finger in use ^25^; surround inhibition is also likely associated with the production of the individuated finger movements ^26^. The biomechanical constraints on the finger dexterity are the anatomical linkages between the tendons and muscles of the hand and forearm ^6,27^. Despite these constraints, highly-skilled movements such as playing the piano are typically characterized by exceptionally independent movements between the fingers, which outperforms those of untrained individuals ^18^. One unanswered question is whether such motor expertise is associated with adaptive changes in the neuromuscular or biomechanical constraints. Neuroplastic adaptations in the sensorimotor system responsible for skillful finger movements have been observed in musicians ^9,28,29^. However, several studies reported reduced surround inhibition of the finger muscles in musicians compared with non-musicians ^26,30^, which may predict an increase in neuromuscular constraints. As for the biomechanical constraints, little has been known about the anatomical characteristics of the fingers of trained musicians.

The primary purpose of the present study is to define the expertise-dependent characteristics of the hand neuromuscular and biomechanical functions in pianists capable of highly individuated finger movements by comparing with musically-untrained individuals. However, even among trained pianists, there are large inter-individual differences in motor dexterity ^31,32^. In addition to deliberate practice ^33^, recent studies demonstrated that innate biological features underlie individual differences in musicians’ motor expertise ^34–36^. This raises the possibility that even neuromuscular and biomechanical features that have no differences between pianists and non-musicians can explain an individual difference in the finger motor dexterity across trained pianists. The secondary purpose of the study is therefore to identify the neuromuscular and biomechanical features of the hand in association with an individual difference in the finger motor dexterity across trained pianists.

## MATERIAL AND METHODS

### Participants

The present study consisted of two studies. In the study 1, 10 right-handed pianists (5 female) and 10 age-matched right-handed non-musicians (5 female) took part in the experiments (Table 1). In the study 2, 14 right-handed pianists (21.1 ± 1.3 years, 9 female) participated. All pianists majored in piano playing and underwent formal musical education at music conservatories, whereas the non-musicians had very little or no experience of piano training (less than 5 years). We also asked the pianists their history of piano training (i.e., the age at which piano training was initiated and their average practice time at each age) in each pianist. In accord with the Declaration of Helsinki, the experimental procedures were explained to all participants. Informed consent was obtained from all participants prior to participation in the experiment. The experimental protocol was approved by the ethics committee at Sophia University, and all methods were performed in accordance with the relevant guidelines and regulations.

**Table 1.**
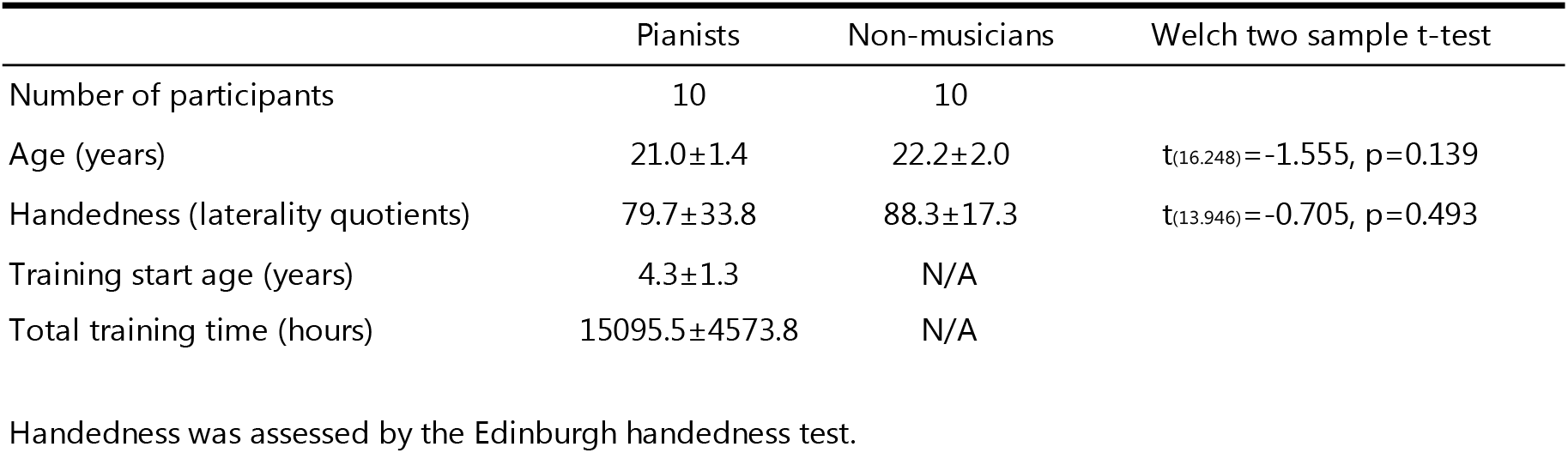
Information on participants of the study 1.

### Experimental Tasks

The experiment consisted of assessments of the neuromuscular and biomechanical functions of the right hand. This included the individuated movements, maximum isometric force production, maximum rate of repetitive movements, and range of motion of each of the individual fingers.

#### Assessment of the individuated finger movements

We investigated active and passive individuated finger movements in each participant. Each participant was seated and put his/her forearm on a table with the shoulder in 20 degree of flexion, and with the elbow flexed to 90 degree. The forearm was pronated and the wrist was in a neutral position. For the active motion condition, each participant voluntarily alternately flexed and extended each of his or her fingers, with a metronome providing a tempo of 184 beat per minute (BPM) (i.e. either flexion or extension per a beat). The instructed range of motion was approximately 60 degrees of the metacarpophalangeal (MCP) joint. The participants were instructed to move without looking at their hand and to keep the remaining fingers unmoved as much as possible ^27^. Five trials were performed for each of the index, middle, ring, and little fingers. For the passive motion condition, each participant wore a custom-made hand exoskeleton that generated force to the proximal-phalange of each finger so as to flex and extend the finger (exiii, Inc.). The exoskeleton moved each finger at the same pace as the active condition (i.e. 184 BPM). In both the active and passive motion conditions, we recorded dynamic changes in the finger joint angles by using sensors embedded in a right-handed custom-made glove (CyberGlove III, CyberGlove Systems Inc.). We recorded the motions at 15 degrees of freedom with angular resolution <0.5°, at 8.3 msec intervals (i.e., sampling frequency = 120 Hz). The measured angles were the metacarpophalangeal (MCP) and proximal phalangeal (PIP) joint angles of the four fingers.

### Maximum force production test

Each participant was asked to exert the maximal force isometrically in each of four different directions (i.e. flexion, extension, leftward, rightward) over three seconds with each of the four fingers. The forearm was pronated (i.e. palm down), and the fingers were at the neutral position. The location of the sensors was adjusted so that the distal phalanx could exert the force to the sensor. We recorded the exerted force using a custom-made force sensor system (Leading-Edge Research and Development Accelerator, Inc.) by 1 kHz ^32^. The resolution and maximum measurable force of the sensor were 0.05 and 49 N, respectively. During the isometric force production by the finger, the participants were instructed to keep the remaining four digits immobilized voluntarily. The wrist was immobilized using Velcro tape so that the wrist movements could not affect the force production. The peak value of the force production was defined as the maximum force.

#### Maximum rate of repetitive finger movements

Each participant repetitively tapped the force sensor as fast as possible for 10 seconds with each of the fingers in each of four different directions (i.e. flexion, extension, leftward, rightward). The arm and hand posture and the location of the sensors were the same as those at the maximum force production test. The wrist was immobilized using Velcro tape so that the wrist movements could not affect the finger motions. The total number of the taps per a second was defined as the maximum rate of repetitive movements by each finger for each direction.

#### Range of motion

The maximum range of motion for the leftward and rightward directions (i.e. lateral motion) of each of the fingers was evaluated in each of active and passive conditions. In the active condition, each participant was asked to move each finger to the left and right maximally while keeping the hand open as much as possible. In the passive condition, the experimenter moved a finger of each participant to the left and right while keeping the hand open as much as possible. In both conditions, the forearm was pronated with the palm facing down. The postural muscular contraction was asked to be minimized. The difference of the angles between the leftward and rightward motions was defined as the range of motion for each finger.

### Data Analysis

Using the finger kinematic data derived from the data-glove during the individuated finger movements, we computed the individuation index (II), active contribution index (ACI), and passive contribution index (PCI), based on the methods established in a previous study ^27^. For each trial, the average angular excursion at each of the MCP and PIP joints of each finger was computed. The II is a measure of the independent movement of a single finger that was instructed to be moved without moving the other fingers. The II was defined as 1 minus the relative joint excursions in the fingers that were instructed to remain immobile (i.e. noninstructed fingers), as follows:

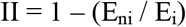

where E_ni_ is the average excursion of both the MCP and PIP joints of the noninstructed fingers, and E_i_ is the average excursion of both the MCP and PIP joints of the instructed finger. The II will be akin to 1 for an ideally individuated movement where the instructed finger moved without any movement of the non-instructed fingers. The closer the II is to 0, the more noninstructed finger movement occurred together with the instructed finger movement.

To assess the effects of biomechanical coupling of the fingers on finger independence, a PCI was computed for each instructed finger according to the joint excursion value in the passive motion condition. Here, the PCI was defined as the average angular excursion of the joints of the noninstructed fingers expressed as a fraction of the average excursion of the MCP and PIP joints of the instructed finger, as follows:

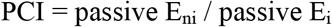

The PCI will be close to 0 with smaller motions of the noninstructed fingers and will increase with increases in the noninstructed finger motions in the passive condition.

To quantify impacts of neuromuscular control on finger independence, an ACI was computed for each of the fingers based on joint excursion values derived from the passive and active conditions. The ACI was defined as the relative excursion of the MCP and PIP joints of the noninstructed fingers in the active motion condition minus the PCI, as follows:

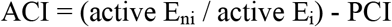

The ACI will be close to 0 with smaller motions of the noninstructed fingers or if they moved the same amount in both the passive and active motion conditions. The ACI will increase with an increase in the noninstructed finger movement in the active motion condition beyond that seen in the passive motion condition.

#### Statistics

The present study evaluated the following variables; II, PCI, ACI, maximum force, maximum tapping rate and its coefficient of variation (CV), and range of motion of each of the index, middle, ring, and little fingers.

In the study 1, a group-wise comparison was performed statistically. For each of the II, PCI, and ACI, two-way mixed-design analysis of variances (ANOVA) with independent variables using finger (4 levels: index, middle, ring, and little fingers) and group (2 levels: pianists and non-musicians) were performed. A Kolmogorov-Smirnov test confirmed a normal distribution of each of all variables evaluated for each group (p > 0.05). For each of the maximum force, maximum tapping rate and its CV of all movement directions (flexion, extension, leftward, and rightward), multiple analysis of variance (MANOVA) using Pillai’s trace statistics were performed. MANOVA was also performed for the range of motion in the active and passive conditions, which used finger and group as independent variables. Post-hoc tests with correction for multiple comparisons ^37^ were performed in the case of significant results of ANOVA and MANOVA. These statistical analyses were performed with R statistical software (“manova” function for MANOVA and “ezPerm” function for ANOVA).

#### Penalized elastic net regression

In the study 2, we tested whether the inter-individual differences in the measure of the dexterity (defined as the maximum tapping rate) of the individual fingers were associated with those in the measures of neuromuscular characteristics of the fingers (i.e. II, PCI, muscular strength, range of motion) across pianists. We performed penalized multiple regression analyses with L1 and L2 norm regularization in a linearly combined fashion, so called elastic net regression ^38^. We used this penalized regression method firstly because it works independently of multicollinearity between the variables and secondly because the analysis is robust even with a limited number of samples. Also, this regression is not a conventional statistical test but machine learning, and thus neither correction for tests with multiple variables and data distribution needs to be concerned. The analysis was performed with R (“glmnet” function). Both the λ parameter that determines the overall intensity of regularization and the α parameter that specifies a ratio between the L1 and L2 regularization were optimized through an iteration of ten-fold cross-validation of the regularized regression model so as to minimize the least-squared error of the model. The independent variables in the regression model included both age at which piano training was commenced and the total amount of life-long piano practicing of the pianists, which were derived from a questionnaire (Table 1).

## RESULTS

### Study 1: Comparison between pianists and non-musicians

Figure 1 illustrated the group means of the II, PCI, and ACI of the index, middle, ring, and little fingers in the pianists and non-musicians. Two-way mixed-design ANOVAs yielded no interaction effect between group and finger (F(3, 54) = 0.90 and 1.47 and p = 0.448 and 0.232 for the II and ACI, respectively) but significant main effects of both group (F(1, 18) = 4.71 and 5.38 and p = 0.044 and 0.032 for the II and ACI, respectively) and finger (F(3, 54) = 23.64 and 7.71 and p = 6.649 × 10^−10^ and 2.223 × 10^−4^ for the II and ACI, respectively). Post-hoc tests identified a significant group difference in the II and ACI only for the middle finger, showing higher movement independence and smaller neuromuscular constraint for this finger for the pianists than the non-musicians. For the PCI, neither an interaction effect (F(3, 54) = 1.59 and p = 0.201) nor a main effect of group (F(1, 18) = 0.484 and p = 0.496) was significant, which confirmed no group difference for any of the fingers.

**Figure 1.**
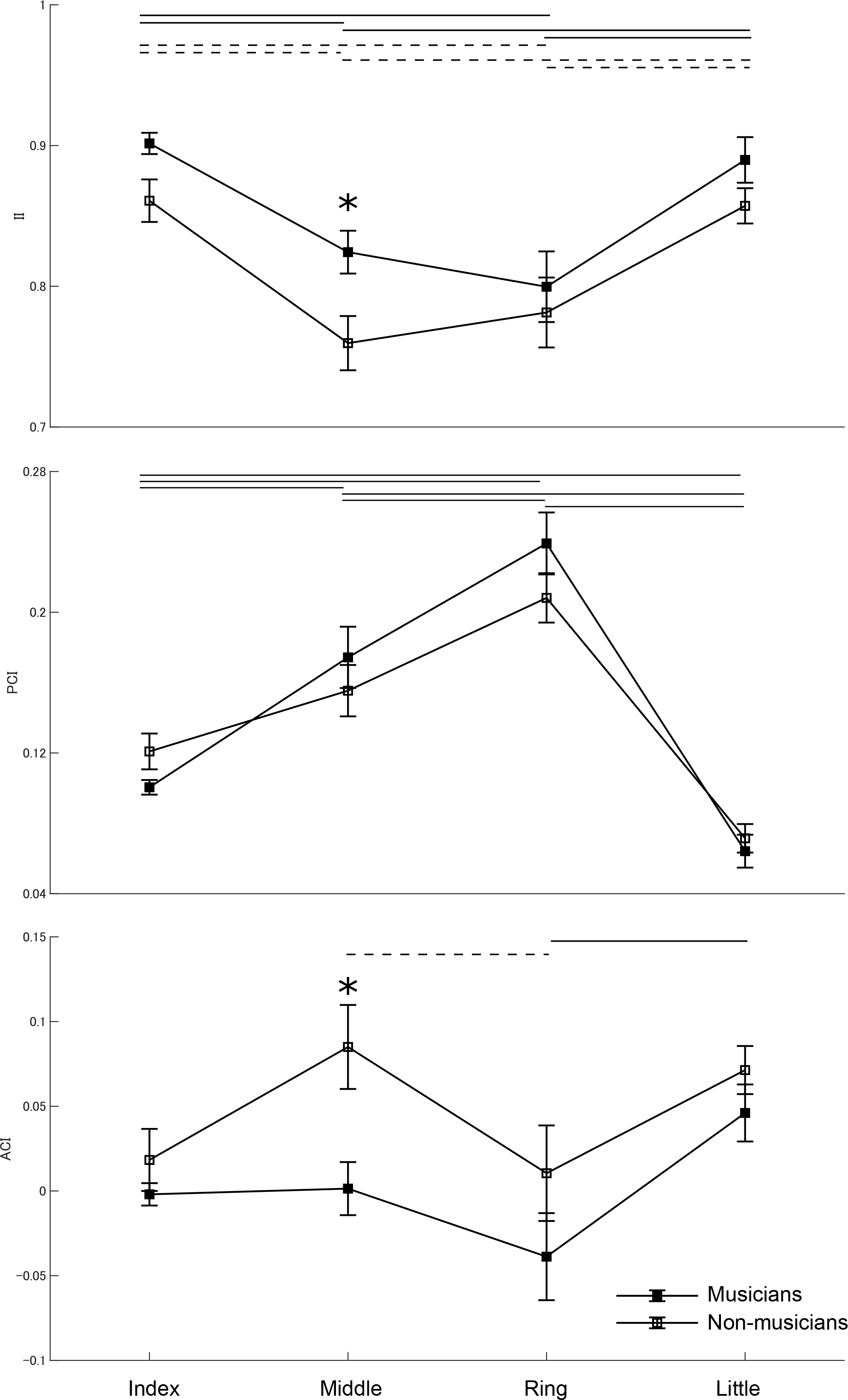
Group means of the individuation index (II), passive contribution index (PCI), and active contribution index (ACI) at the index (I), middle (M), ring (R), and little (L) fingers in the pianists (filled square) and non-musicians (open square). A solid and dotted horizontal line indicates a significant difference between the fingers and between the pianists and non-musicians, respectively (p < 0.05). *: significant group difference (p < 0.05). The higher value of the II and ACI indicates higher movement individuation and neuromuscular constraints on the fingers, respectively. An error bar indicates one SEM.

Figure 2 shows the group means of the maximum force exerted in four different directions and the range of lateral motion of the fingers at the active and passive conditions in the pianists and non-musicians. For the maximum force, MANOVA showed that neither interaction effect between group and finger (F(4,16) = 0.52, p = 0.937) nor group effect (F(1,4) = 2.33, p = 0.062) was significant. For the range of motion, although MANOVA revealed a significant group effect (F(1, 2) = 5.32, p = 0.007), ANOVAs did not yield any significant group effect at the active (F(1, 2) = 1.03, p = 0.310) and passive (F(1, 2) = 2.50, p = 0.120) conditions. These findings confirmed no group difference in the finger muscular force and range of motion for any of the fingers between the pianists and non-musicians. MANOVA also showed a significant finger effect on the muscular force (F(4, 16) = 3.30, p = 0.211 × 10^−4^) and ROM (F(3, 6) = 12.09, p = 0.558 × 10^−10^).

**Figure 2.**
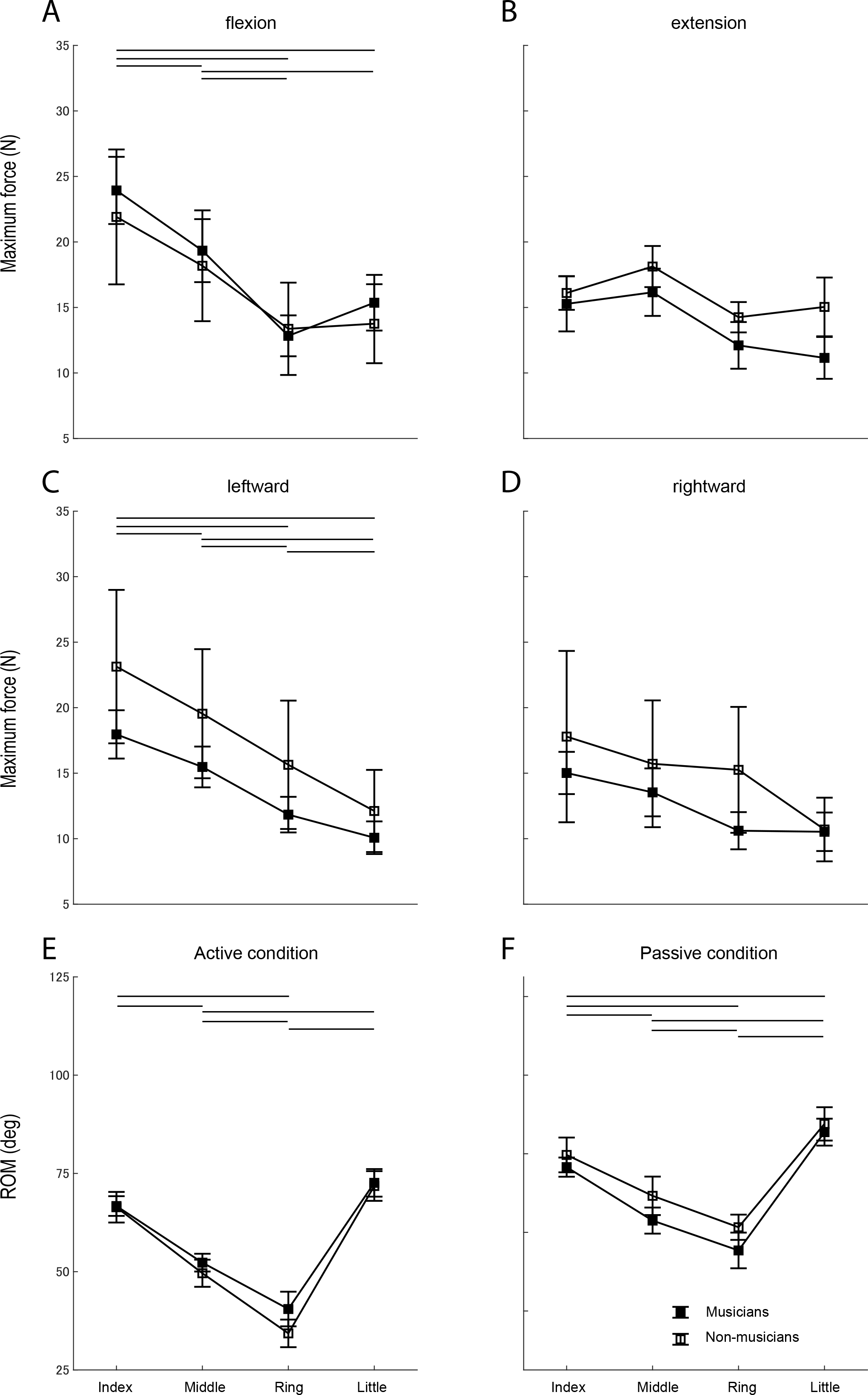
A-D: Group means of the maximum force exerted by the index (I), middle (M), ring (R), and little (L) fingers in each of the flexion, extension, leftward, and rightward directions in the pianists (filled square) and non-musicians (open square). The leftward and rightward directions indicate when the hand is pronated so that the palm faces down. E-F: Group means of the range of lateral motion of each finger at the active (E) and passive (F) conditions in the pianists (filled square) and non-musicians (open square). None of these variables showed a significant group difference. A solid horizontal line indicates a significant difference between the fingers in all participants pooled (p < 0.05).

Figure 3 displays the group means of the maximum tapping rate and CV of the inter-tap intervals for each of the fingers in the pianists and non-musicians. For the maximum tapping rate, MANOVA yielded a significant group effect (F(1, 3) = 15.75, p = 6.321 × 10^−8^), but neither group and finger interaction nor finger effects was significant. Two-way mixed-design ANOVAs also demonstrated significant effects of group for both the flexion-extension (F(1, 3) = 44.63, p = 4.269 × 10^−9^) and the rightward motion (F(1, 3) = 6.43, p = 0.013) but not for the leftward motion (F(1, 3) = 3.10, p = 0.082). Post-hoc tests identified group differences at the middle, ring, and little fingers for the flexion-extension, and at the ring finger for the rightward motion. For the CV of the inter-tap intervals, MANOVA yielded a significant group effect (F(1, 3) = 9.87, p = 1.65 × 10^−5^) but neither interaction effect of group and finger nor main effect of finger was significant. Two-way mixed-design ANOVAs found a significant interaction effect only for the flexion-extension direction (F(3, 9) = 3.61, p = 0.017), and also a significant group effect for each of the flexion-extension (F(1, 3) = 21.52, p = 1.53 × 10^−5^), rightward (F(1, 3) = 11.42, p = 0.001), and leftward motions (F(1, 3) = 16.86, p = 1.05 × 10^−4^). Post-hoc tests found a significant group difference only at the ring finger during the flexion-extension tapping. These findings confirmed that faster tapping in the pianists was not attributed to a tradeoff between speed and accuracy. We therefore defined the maximum finger tapping rate as a measure of the finger dexterity, which was used in the regression model of the study 2.

**Figure 3.**
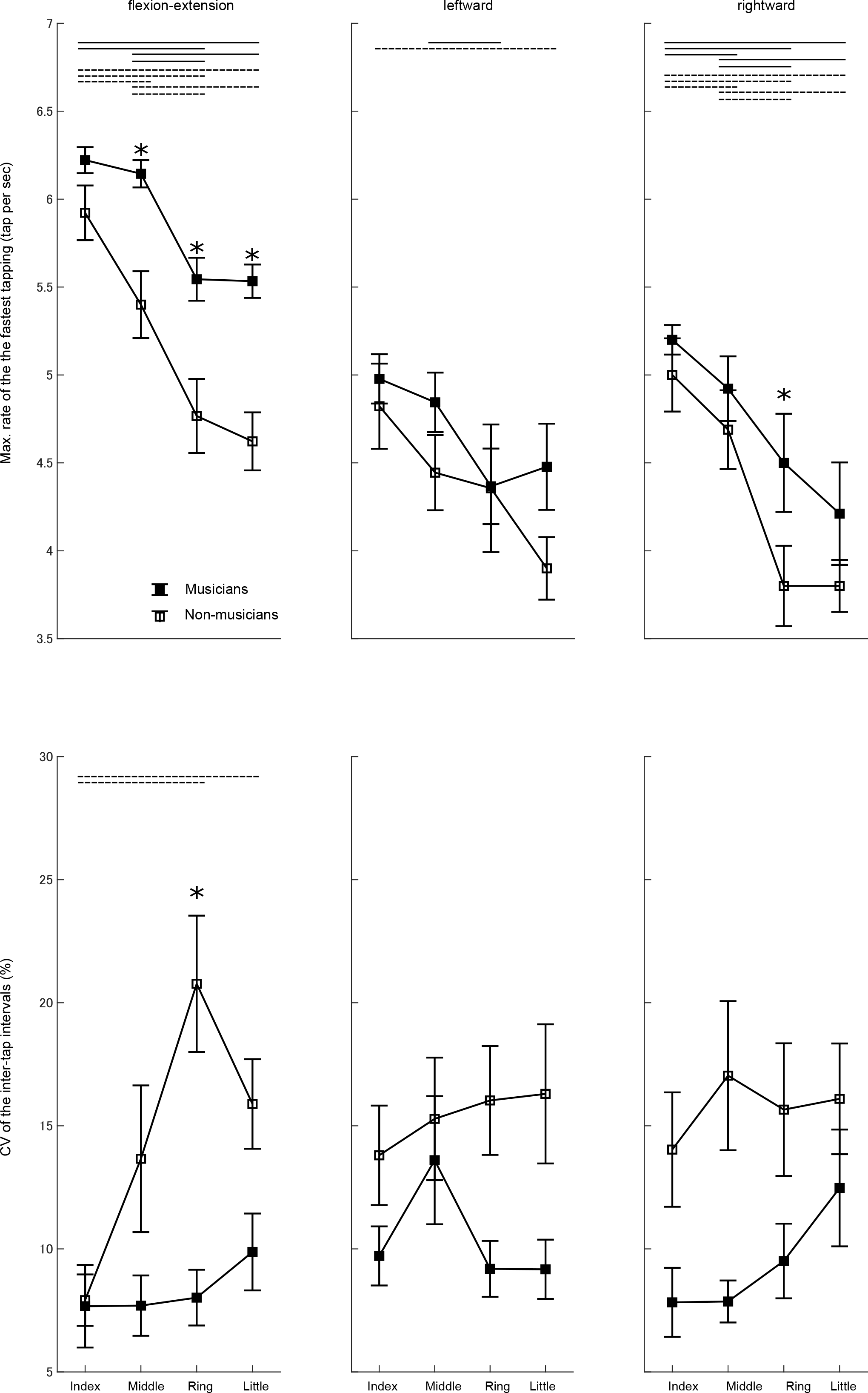
Group means of the maximum rate of the fastest tapping (upper panel) and coefficient of variation (CV) of the inter-tap interval during the fastest tapping (lower panel) with each of the index, middle, ring, and little fingers in the flexion-extension (right panel), leftward (middle panel), and rightward (left panel) directions in the pianists (filled square) and non-musicians (open square). A solid and dotted horizontal line indicates a significant difference between the fingers and between the pianists and non-musicians, respectively (p < 0.05). *: significant group difference (p < 0.05). An error bar indicates one SEM.

### Study 2: Individual differences in the maximum movement rate between expert pianists

Table 2 and Figure 4 summarizes the results of the elastic net regression for the maximum finger tapping rate in the pianists who participated in the study 2. The regression selected some neuromuscular features of the fingers associated with the maximum movement rate of each finger. The maximum tapping rate of the middle finger was associated negatively with the PCI of the index finger and positively with the maximum flexion force of the ring finger. The maximum tapping rate of the ring finger covaried positively with the maximum flexion force of the index finger. The maximum tapping rate of the little finger was associated positively with the maximum flexion force of the index finger and negatively with the range of motion of the middle finger at the passive condition. Importantly, the elastic net demonstrated that none of the maximum rates of the fastest finger-tapping covaried with age of starting piano training and the total amount of piano practicing. In summary, these findings indicate that the maximum rate of the fastest finger tapping of the expert pianists was associated with the biomechanical features of the hand (i.e. passive range of motion and PCI) and maximum finger muscular force, both of which displayed no significant group difference in the study 1. In addition, none of these predictors had a significant correlation with age of starting piano training and the total amount of piano practicing (p > 0.05).

**Table 2.** Results of elastic net multiple regression analyses for the fast individual finger movements of pianists.

| Dependent variables | max. tap rate 3 |  | max. tap rate 4 |  | max. tap rate 5 |  |
| --- | --- | --- | --- | --- | --- | --- |
| Independent variables | PCI2 | $-2.41 \times 10^{-1}$ | MVC Flexion2 | $3.30 \times 10^{-1}$ | MVC Flexion2 | $2.77 \times 10^{-1}$ |
| | MVC Flexion4 | $3.90 \times 10^{-2}$ | | | Passive ROM3 | $-1.97 \times 10^{-2}$ |
| Alpha |  | 0.99 |  | 0.99 |  | 0.99 |
| Lambda |  | 0.33 |  | 0.35 |  | 0.36 |

MVC: maximum voluntary contraction
ROM: range of motion
2, 3, 4, 5 in the variables indicates the index, middle, ring, and little finger, respectively.

**Figure 4.**
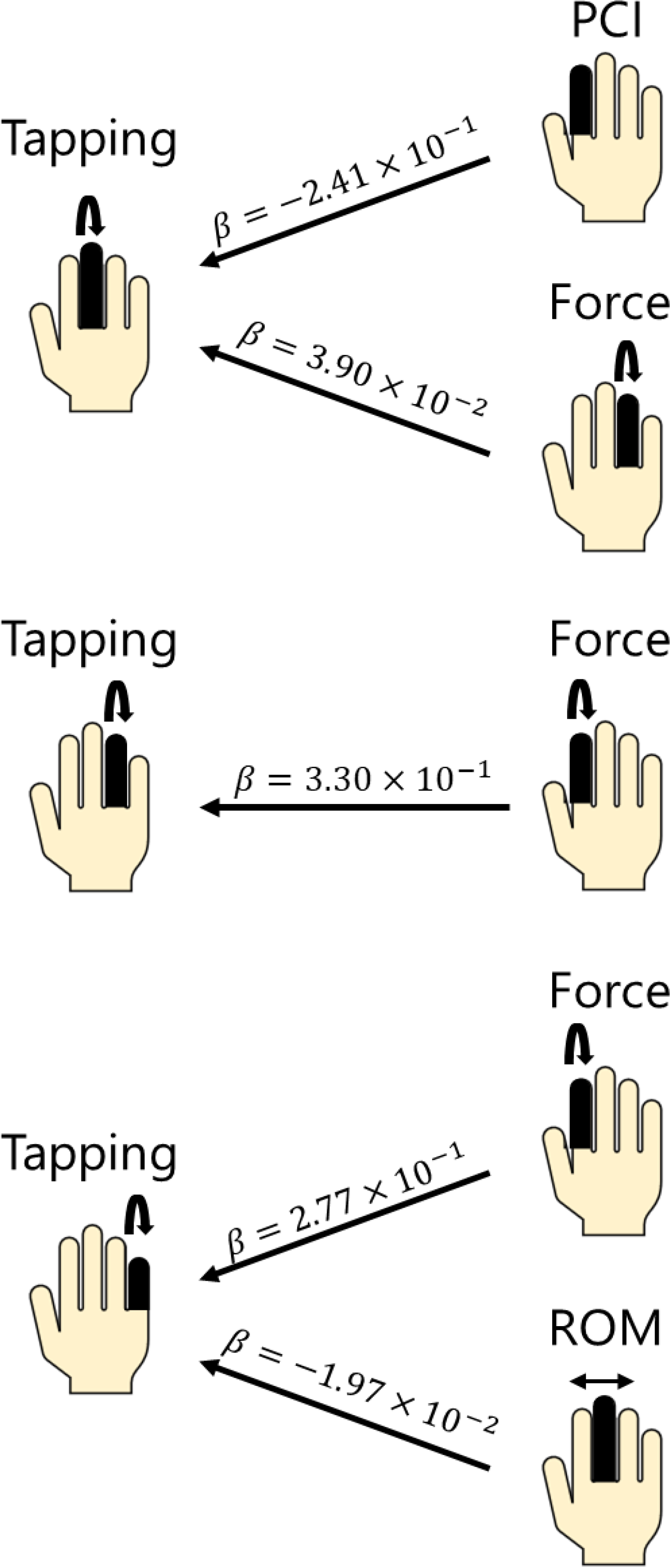
Visualized results of the elastic net regression (Table 2). Variables accounting for the maximum rate of the fastest tapping with each of the middle, ring, and little finger are displayed at the right column. The beta value indicates the coefficient derived from the regression.

## DISCUSSION

The present results consist of two major observations. First, compared with non-musicians, pianists exhibited faster and more individuated movements of the fingers. In addition to smaller ACI in the pianists, a lack of any group differences in PCI, the range of motion, and the muscular strength provided evidences of neuromuscular but not biomechanical adaptation of the fingers through extensive piano training. Second, an individual difference in the maximum movement speed of the individual fingers between the trained pianists covaried with the range of motion, PCI, and muscular strength of the fingers, but not with movement independence between the fingers and ACI. The results indicate that both the muscular strength and a measure of mechanical coupling between the fingers, which had no group difference, account for the individual difference in the finger dexterity across pianists. Importantly, none of these finger characteristics was correlated with early and deliberate piano practicing, which includes age of starting to play the piano and total duration of piano practicing. Importantly, the contrasting observations between these two results highlight distinct differences in biological factors associated with the effects of extensive piano training on and between-pianists differences in the finger motor dexterity. Extensive piano training is likely to reduce neuromuscular constraints of the fingers so as to improve the independent movement control, but not necessarily ensure mastery of the finger motor virtuosity of the experts due to the biomechanical constraints and limited muscular strength.

The superior independence of movements between the fingers in pianists to that of non-musicians accompanied reduced neuromuscular constraints of the fingers without differences in biomechanical constraints. This finding extends the previous behavioral studies that demonstrated enhanced movement individuation between the fingers through musical training ^18,39^. Some neurophysiological adaptation can be proposed as a putative mechanism underlying the reduced neuromuscular constraints. Using transcranial magnetic stimulation, several studies demonstrated neuroplastic adaptation of the motor cortical functions responsible for dexterous finger movements through musical training ^9,26,30,40^. For example, patterns of movement coordination between the fingers, which are encoded in the motor cortex, differed between the pianists and non-musicians ^9^. Motor cortical adaptation caused by musical training also includes reduced surround inhibition in musicians, which strengthens coupling between multiple finger muscles ^26^. This suggests that the enhanced movement individuation between the fingers result not from enhanced functional individuation between the adjacent finger muscles, but from neuroplastic alteration of finger muscular coordination so that force production at a finger muscle accompanies the production of adjacent-fingers’ muscular force counteracting against neuromuscular and mechanical coupling between the fingers.

A key finding of our study was the association of the individual differences in the finger motor dexterity between the pianists with mechanical coupling between the fingers and muscular force strength but not with early and deliberate musical training nor with neuromuscular factors of the finger motor independence. A lack of any relationship of the finger dexterity of the pianists with deliberate musical practice corroborates a concept of gene-environment interaction in acquisition of motor virtuosity ^35,36,41^. Possible hereditary factors include innate individual differences in physiological properties of muscles ^42,43^ and hand anatomy ^44^. In addition, the way of practicing can also play a role in mastering the finger motor skills ^34^. For example, the effects of piano practicing on the finger movement independence depend on explicit attention to movement accuracy ^39^. The acquired movement coordination between the fingers through practicing the piano also differs between implicit and explicit motor learning ^40^. Furthermore, training and exercise being performed independently of musical practicing, such as muscle strength training and stretching to open the hand, may also aid in sophisticating finger motor skills, on the basis of our finding that the finger muscular strength and inter-finger mechanical coupling covaried positively with the maximum speed of the finger movements, although our results are correlational and any causal relationships are not yet demonstrated. Although a causal relationship of these finger characteristics with heredity, ways of musical practicing, and training and exercising independent of musical practicing remains unknown, the present findings can help in optimizing training and practicing of acquiring and sophisticating finger motor skills. Due to a large number of potential biological factors associated with the finger dexterity, it is difficult to specify a target for motor training and rehabilitation. This can cause that even through long-term extensive training, experts are incapable of mastering motor skills, which often leads to overtraining and thereby triggers maladaptive changes in the sensorimotor system such as musicians’ dystonia ^11^. Our findings and methodologies successfully identified a small number of neuromuscular and biomechanical factors associated with the finger dexterity, which will potentially enable us to focus on a target for motor training and musical practicing for optimizing the finger motor dexterity while reducing risks of overuse syndromes.

## Acknowledgements

We thank Mr. Ryuya Tanibuchi for his assistance in developing scripts for controlling the exoskeleton, and Dr. Masato Hirano for his constructive suggestions to this research. This study was supported by JST CREST.

## AUTHORSHIP

S.F. and Y.K. developed the study concept and design. Y.K. performed the data collection, data analyses, and statistics. S.F. and Y.K. interpreted the data and drafted the manuscript. S.F. and Y.K. approved the final version of the paper for submission.

